## Supplementary material for "Rational engineering of an elevator-type metal transporter ZIP8 reveals a conditional selectivity filter critically involved in determining substrate specificity": SI

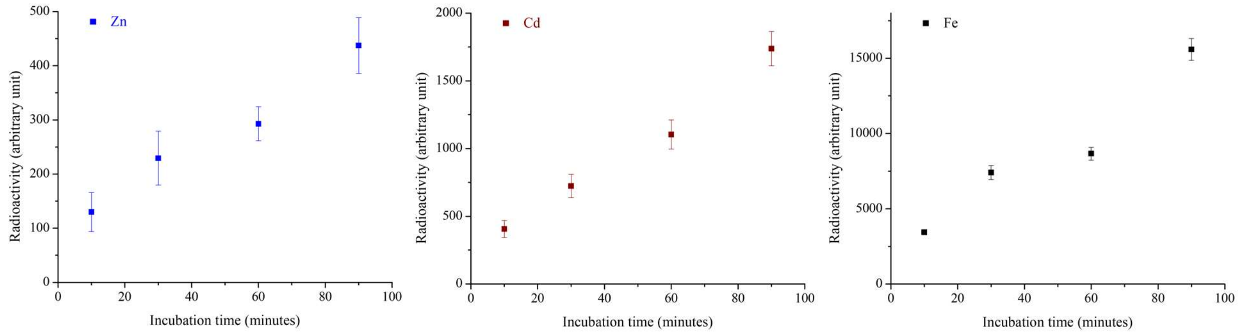


**Figure S1.** Time courses of metal transport by the wild type ZIP8. *left*: $Zn^{2+}$; m*iddle*: $Cd^{2+}$; *right*: $Fe^{2+}$. Metal substrates were detected using a gamma counter (for $^{65}Zn$ and $^{109}Cd$) or a liquid scintillation counter (for $^{55}Fe$). The concentrations of $Zn^{2+}$, $Cd^{2+}$, and $Fe^{2+}$ were 5 μM, 5 μM, and 20 μM respectively. The shown data are from one representative experiment of three independent experiments with three replicates for each condition. The bars indicate S.D.

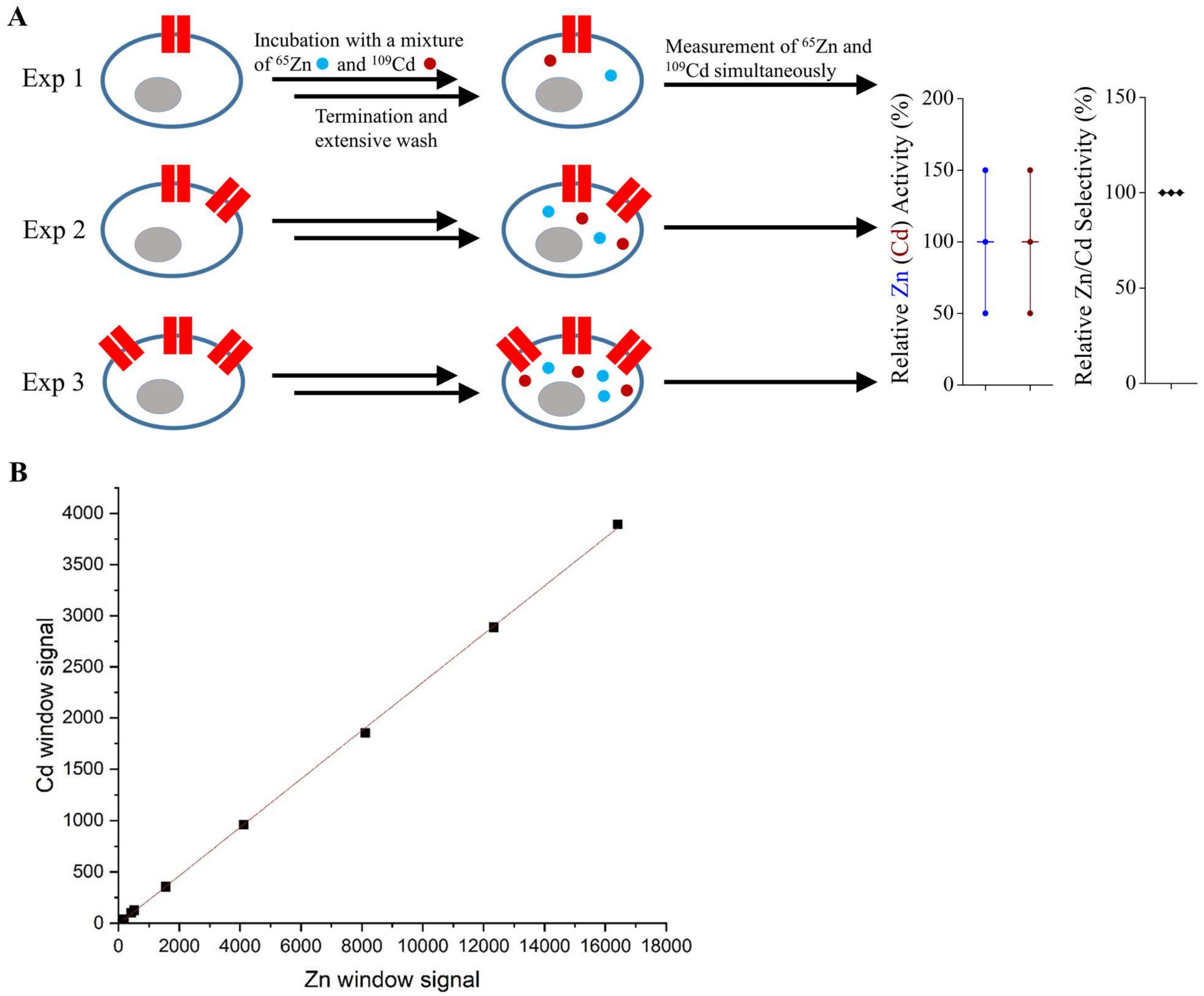


**Figure S2.** Internal competition transport assay and data processing. (**A**) Illustration of the experimental procedure. Cells expressing different levels of ZIP8 (shown as red channels) in three independent experiments (Exp 1-3) are incubated with a mixture of $^{65}Zn$ and $^{109}Cd$ under the same experimental conditions, followed by termination with an ice cold EDTA-containing solution and extensive wash. Radioactivities of $^{65}Zn$ and $^{109}Cd$ associated with the cells were simultaneously quantified by using a gamma counter in two detection windows (800-1500keV for Zn and 30-150keV for Cd). As demonstrated in the simulated experiments, although the Zn (or Cd) transport activity may have a large deviation due to varied expression levels of the transporter in different experiments, the ratios of the transport activities of Zn to Cd (the Zn/Cd selectivity) are essentially the same. (**B**) Calibration of the signals recorded in the Cd window. Radioactivities of a series of $^{65}Zn$ standard samples (0-20 μM) recorded in the Cd window are plotted against the readings recorded in the Zn window. The slope of the curve (0.235) means that, for any given sample containing $^{65}Zn$ and $^{109}Cd$, 23.5% of the reading recorded in the Zn window contributes to the reading in the Cd-window. Therefore, to calibrate the readings in the Cd window, 23.5% of the reading in the Zn window was subtracted from the reading recorded in the Cd window to determine the radioactivity truly derived from $^{109}Cd$. $^{109}Cd$ does not contribute to the reading in the Zn window.

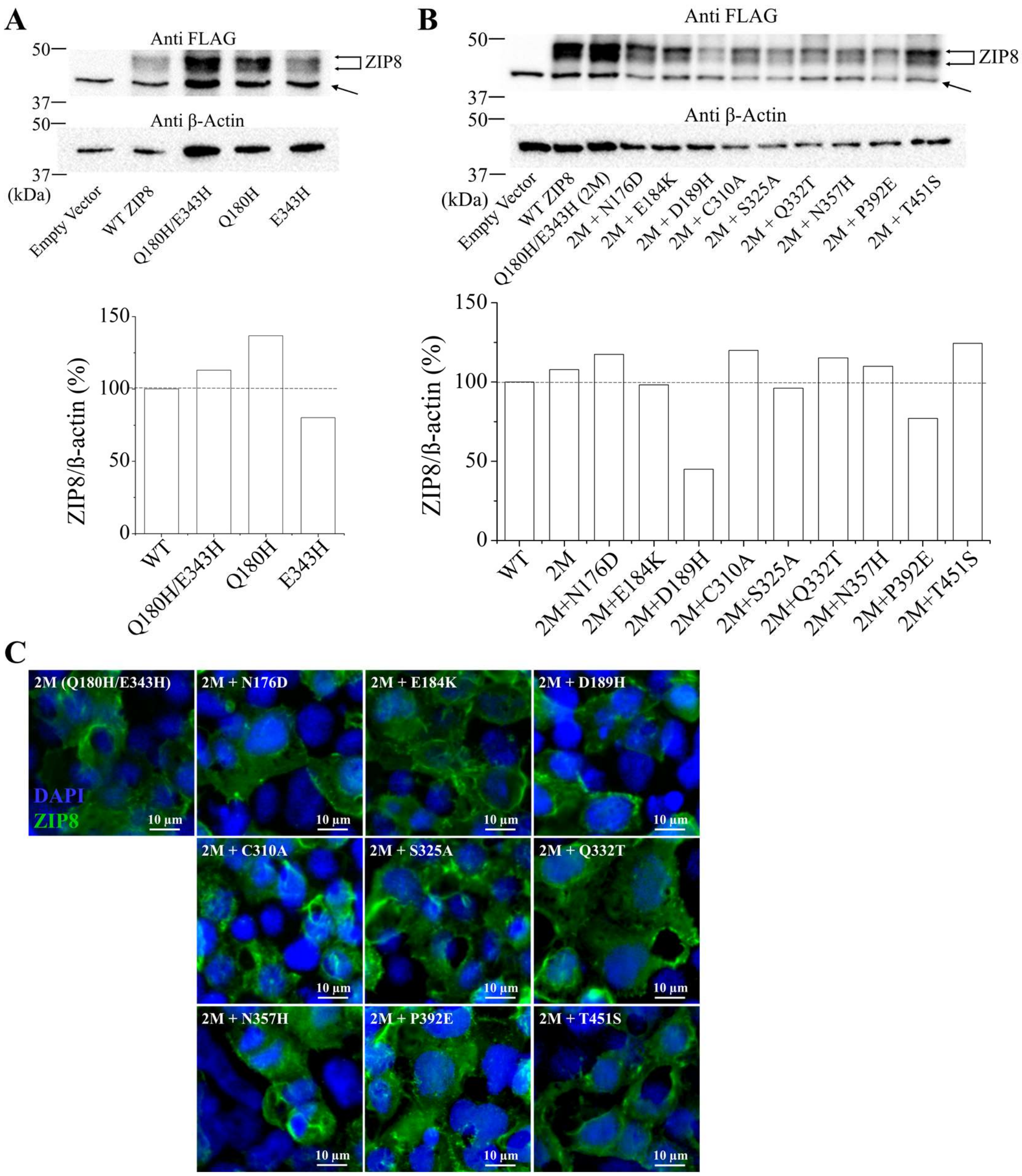


**Figure S3.** Expression analysis of ZIP8 and the variants. (**A**) Comparison of the single and double variants with the wild type ZIP8 by Western blot (upper panel) and quantification analysis using ImageJ (lower panel). (**B**) Comparison of triple variants with the wild type ZIP8 and the Q180H/E343H (2M) variant by Western blot (upper panel) and quantification analysis using ImageJ (lower panel). N-FLAG ZIP8 and β-actin were detected using anti-FLAG and anti-actin antibodies, respectively. The expression level of a variant is expressed as the percentage of the ratio of the wild type ZIP8 over β-actin. The shown result is from a representative experiment. (**C**) Immunofluorescence analysis of cell surface expressed ZIP8 variants. The procedure is the same as indicated in the legend of Figure 4A.

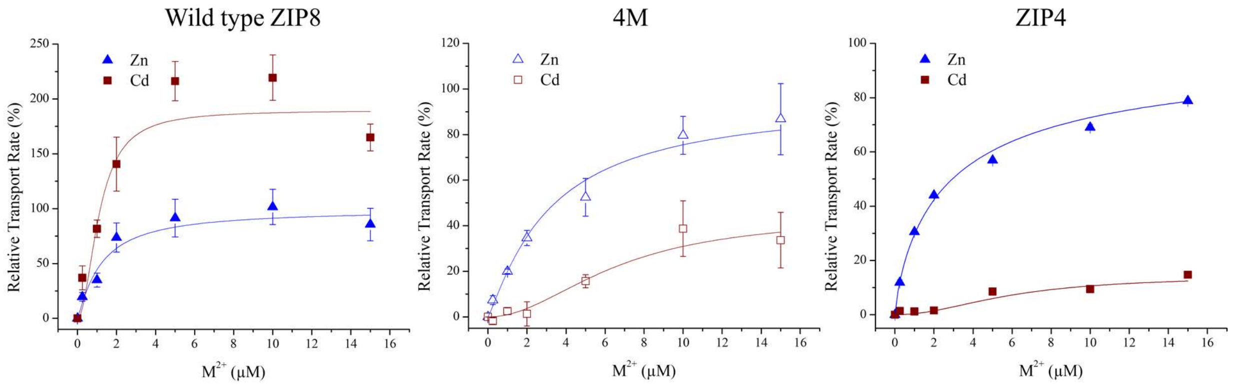


**Figure S4.** The Zn and Cd transport activities of the wild type ZIP8 and the 4M variant (*left*) in comparison with human ZIP4 (*right*). The same data shown in Figure 3B are presented differently for better comparison with ZIP4. The transport rates are expressed as the percentages of the $V_{max}$ for $Zn^{2+}$. The shown data are from one representative experiment of three independent experiments with three replicates for each condition. The bars indicate S.D.

**A**

| Metal ion | Imidazole (His) | | | Acetate (Asp and Glu) | | |
|---|---|---|---|---|---|---|
| | logK | Energy (kcal/mol) | Polarizability on coordinating nitrogen | logK | Energy (kcal/mol) | Polarizability on coordinating oxygen |
| $Zn^{2+}$ | 2.55 | -3.48 | 2.71 | 1.58 | -2.16 | 0.77 |
| $Fe^{2+}$ | 1.80 | -2.46 | N/A | 1.40 | -1.91 | N/A |
| $Mn^{2+}$ | 1.20 | -1.64 | N/A | 1.40 | -1.91 | N/A |
| $Cd^{2+}$ | 2.66 | -3.63 | 3.20 | 1.93 | -2.63 | 1.17 |

**B**

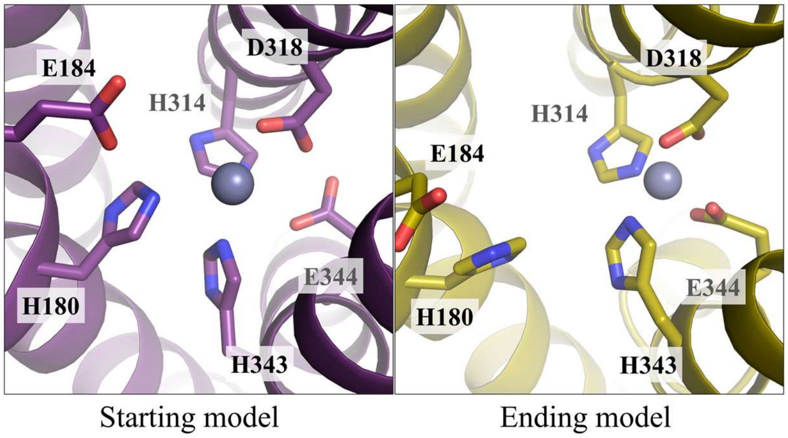


**Figure S5.** Computational characterization of metal binding at the selectivity filter. (**A**) Free energy changes of metal ion binding with small molecule ligands (refs 41&42). (**B**) The initial trial of MD simulation. The zinc ion initially located at the selectivity filter (*left*) moved to the transport site in the ending model (*right*) where it is coordinated with the residues from M1 (H314, H343), M2 (E344), and a bridging residue involved in both M1 and M2 (D318).

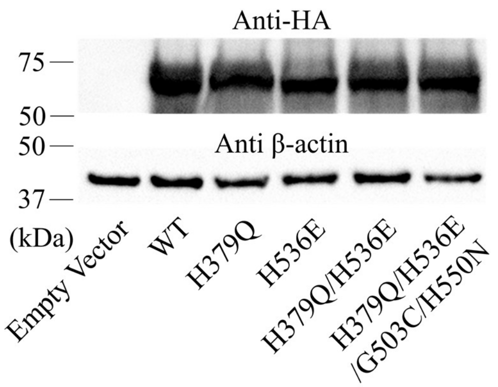


**Figure S6.** Expression analysis of ZIP4 and the variants by Western blot. Human ZIP4 with a C-terminal HA tag and β-actin were detected using anti-HA and anti-β-actin antibodies, respectively.

**Table S1.** Sequence identity of the human LIV-1 proteins

| (%) | ZIP7 | ZIP13 | ZIP4 | ZIP12 | ZIP8 | ZIP14 | ZIP5 | ZIP6 | ZIP10 |
|---|---|---|---|---|---|---|---|---|---|
| ZIP7 | 100 | 35 | 27 | 23 | 22 | 24 | 28 | 25 | 27 |
| ZIP13 | 35 | 100 | 29 | 24 | 21 | 24 | 29 | 23 | 23 |
| ZIP4 | 27 | 29 | 100 | 32 | 31 | 30 | 31 | 26 | 25 |
| ZIP12 | 23 | 24 | 32 | 100 | 29 | 31 | 26 | 25 | 27 |
| ZIP8 | 22 | 21 | 31 | 29 | 100 | 48 | 27 | 31 | 30 |
| ZIP14 | 24 | 24 | 30 | 31 | 48 | 100 | 25 | 30 | 29 |
| ZIP5 | 28 | 29 | 31 | 26 | 27 | 25 | 100 | 32 | 35 |
| ZIP6 | 25 | 23 | 26 | 25 | 31 | 30 | 32 | 100 | 40 |
| ZIP10 | 27 | 23 | 25 | 27 | 30 | 29 | 35 | 40 | 100 |

**Table S2.** The primers for mutagenesis.

| Name | Sequence (5’-3’) [a] |
|---|---|
| **ZIP8** | |
| E343H | ACTTCCATAGCAATCCTATGTCACGAGTTTCCCCACGAGTTAGGA |
| C374L | CTATTCAACTTCCTTTCTGCATTGTCCTGCTATGTTGGGCTAGCT |
| C374L/C376A | CTATTCAACTTCCTTTCTGCATTGTCCGCTTATGTTGGGCTAGCTTTTGGC |
| Q180H | CTTTTTTCAAATGCAATTTTCCACCTTATTCCAGAGGCATTTGGA |
| Q180H/E343H/N176D | GCTATTGGGACTCTTTTTTCAGATGCAATTTTCCACCTTATTCCA |
| Q180H/E343H/E184K | GCAATTTTCCACCTTATTCCAAAGGCATTTGGATTTGATCCCAAA |
| Q180H/E343H/D189H | ATTCCAGAGGCATTTGGATTTCACCCCAAAGTCGACAGTTATGTT |
| Q180H/E343H/C310A | ATTGCCTGGATGATAACGCTCGCTGATGCCCTCCACAATTTCATC |
| Q180H/E343H/S325A | GATGGCCTGGCGATTGGGGCTGCTTGCACCTTGTCTCTCCTTCAG |
| Q180H/E343H/Q332T | TCCTGCACCTTGTCTCTCCTTACAGGACTCAGTACTTCCATAGCA |
| Q180H/E343H/N357H | GGAGACTTTGTGATCCTACTCCACGCAGGGATGAGCACTCGACAA |
| Q180H/E343H/P392E | TTGGTGGGCAACAATTTCGCTGAGAATATTATATTTGCACTTGCT |
| Q180H/E343H//T451S | ACTGGAAGAAAAACCGATTTCTCATTCTTCATGATTCAGAATGCT |
| **ZIP4** | |
| H379Q | CTCACTGGGGACGCTGTCCTGCAACTGACGCCCAAGGTGCTGGGG |
| H536E | ACCTCGCTGGCCGTGTTCTGCGAAGAGTTGCCACACGAGCTGGGG |
| G503C | CTGCCCTATATGATCACTCTGTGTGACGCCGTGCACAACTTCGCC |
| H550N | GGGGACTTCGCCGCCTTGCTGAACGCGGGGCTGTCCGTGCGCCAA |

[a] Only the forward primers are shown. The reverse primers are reversely complimentary to the sequences of the forward primers.

[b] The Q180H/E343H/C310A/N357H variant was generated by using the primers for the Q180H/E343H/C310A and N357H variants.

**Table S3.** ICP-MS analysis of some *d*-block metals in the culture media (DMEM+10% FBS) before and after the treatment with the Chelex-100 resin.

| (μM) | Mn | Fe | Zn |
|---|---|---|---|
| Before | 0.059 | 3.988 | 3.637 |
| After | 0.043 | 3.702 | 0.091 |
